# A rapidly prototyped, simple yet versatile dynamic breathing Exposure-on-a-Chip for investigating nanoparticle deposition in the alveoli

**DOI:** 10.1101/2025.03.07.642100

**Authors:** Xiangxu Liu, Melih Engur, Chunyu Yan, Felix Dobree, Taima Najjar, Camille Chassat, Yangyumeng Chen, Marc Stettler, Joseph Xavier, Jorge Bernardino de la Serna

## Abstract

We developed and characterized a three-layer microfluidic Exposure-on-a-Chip (EOC) model to replicate the alveolar microenvironment and simulate breathing motions, offering a physiologically relevant platform for studying inhaled nanomedicines. The EOC chip, fabricated from biocompatible polydimethylsiloxane (PDMS), features a fluidic chamber for cell culture, a pneumatic chamber for pressure application, and a thin PDMS membrane (150 µm) separating the chambers to support cell growth and enable mechanical stretching. Xurography and 3D printing were validated as efficient and reproducible fabrication methods. Mechanical characterization, using fluorescent bead tracing, confirmed that the PDMS membrane accurately mimics alveolar breathing motions under physiological conditions (1-8% strain). Biological validation showed that alveolar epithelial and endothelial cells cultured on the EOC formed functional monolayers, maintaining barrier integrity under cyclic and static stretching, separately. To study nanoparticle behavior, we examined the deposition of nanoparticles under dynamic stretching versus static conditions. Significantly fewer nanoparticles accumulated in cells under continuous dynamic exposure with stretching compared to static culture, highlighting the critical role of mechanical forces in nanoparticle-cell interactions. The EOC platform provides a robust and scalable tool for evaluating nanomedicine efficacy in dynamic environments, representing a significant advancement in alveolus-on-chip technology.

## 1 Introduction

Inhaled drug delivery holds significant promise in respiratory nanomedicine, offering targeted lung delivery to enhance therapeutic efficacy while minimizing systemic side effects. ^1–3^ However, evaluating the efficacy and safety of these therapies remains challenging. Traditional models, such as animal studies, are expensive, time-consuming, and fail to replicate key human-specific physiological and pathological conditions. ^4^ Meanwhile, *in vitro* assessments using primary human cell lines often expose cells to nanoparticles under static conditions, inadequately mimicking the complex behavior of inhaled nanoparticles *in vivo*. ^5,6^ These limitations highlight the need for advanced models capable of replicating the intricate three-dimensional (3D) architecture, dynamic movements, and air-liquid interface of lung alveoli. ^7^

The emergence of organ-on-a-chip (OOC) technology offers a transformative approach to addressing these challenges. ^8^ By integrating the latest advancements in biology, fluid dynamics, and microengineering, OOC systems recreate tissue-specific functions in dynamic, biomimetic microenvironments. ^9^ Recently, OOC has gained significant attention for applications in drug screening and toxicity assessment. Compared to animal models and conventional 2D *in vitro* systems, OOC provides a dynamic, biomimetic microenvironment with microscale channels that guide and manipulate fluid flow across cultured human cell lines. ^10,11^

Over the last two decades, OOC studies on alveoli have successfully reconstituted the alveolar-capillary barrier, demonstrating significant potential in predicting physiological responses and injuries that closely mimic *in vivo* conditions. ^12–14^ However, current OOC systems often neglect the rhythmical expansion and contraction characteristic of alveoli during respiration, limiting their physiological relevance for inhaled drug studies. ^15,16^ As the smallest structure of the respiratory system, alveoli play a vital role in facilitating gas exchange during rhythmic respiration while maintaining a continuous barrier to regulate fluid balance in the lung airspace. Recent research has highlighted the importance of mimicking the rhythmic out-of-plane stretching of alveoli, as it better replicates the physiological surface curvature changes associated with alveolar expansion compared to in-plane stretching. ^17,18^ Studies have shown that periodic mechanical stretching can influence lung surfactant tensioactive properties, inhibit alveolar epithelial cell proliferation and promote extracellular matrix secretion. ^19–21^ It can also enhance the nanoparticle translocation and potentially trigger inflammatory responses. ^22^

To address this gap, we developed a multi-functional biomimetic breathing exposure-on-a-chip (EOC) system, incorporating continuous flow and cyclic stretching to replicate the sinusoidal breathing patterns of human alveoli. The multi-layered polydimethylsiloxane (PDMS) design enables the inclusion of airflow and fluid infusion, mimicking the mechanical stress and shear forces experienced by alveolar epithelial and endothelial cells *in vivo*. The components of the extracellular matrix (ECM) were investigated to establish the mechanical support for examining cell behavior and function. By simulating realistic breathing motions, this aims to be a more predictive model of nanoparticle behavior within the respiratory system.

Achieving such precision requires robust and cost-effective fabrication techniques. Photolithography, a widely used method in microfluidic chip mould fabrication, offers high precision but relies heavily on complex and often costly processes. ^23^ Recent advancements, such as xurography and stereolithographic (SLA) 3D printing, have emerged as promising alternatives for rapid prototyping. While xurography is economical and efficient, it is constrained by material limitations (e.g., substrate thickness for deep channels) and design complexity. ^24,25^ Conversely, 3D printing benefits from superior resolution but faces biocompatibility challenges due to residual resin, necessitating meticulous post-processing to ensure compatibility with biological applications. ^26,27^ PDMS, as the most commonly used material for organ-on-chip devices, offers significant advantages due to its biocompatibility, optical transparency, and gas permeability, which are highly favorable for supporting long-term cell growth and monitoring. Notably, its elasticity further facilitates the simulation of mechanical stimulation of cells. *in vitro*. ^9^

In this study, we utilized our three-layered PDMS-based EOC system to investigate inhaled nanoparticle deposition and interactions under physiologically relevant conditions. Human alveolar epithelial lentivirus-immortalized (hAELVi) cells, mimicking alveolar epithelial type *I* cells, and human umbilical vein endothelial cells (HUVECs) were cultured in the system separately to characterize the biological ability of this chip. Both the alveolar epithelial cell and endothelial cell monolayers maintained their barrier function under cyclic pressure, applied to mimic the breathing motion. Dynamic evolution of nanoparticles exposure on cells under breathing patterns offers a rapid and cost-effective method for *in vitro* nanomedicine studies.

## 2 Material and Methodology

### 2.1 Fabrication of the microfluidic chip

The mould fabrication for the EOC was accomplished using the xurography and 3D printing method (Fig.1A). The EOC comprises a fluidic layer and a pneumatic actuation layer, separated by a thin PDMS membrane as shown in Fig.S1A&C. Consequently, two moulds were created: one for the fluidic channel and another for the pneumatic actuation channel. Both Xurography and 3D printing technique were utilized to prepare the moulds.

**Fig. 1.**
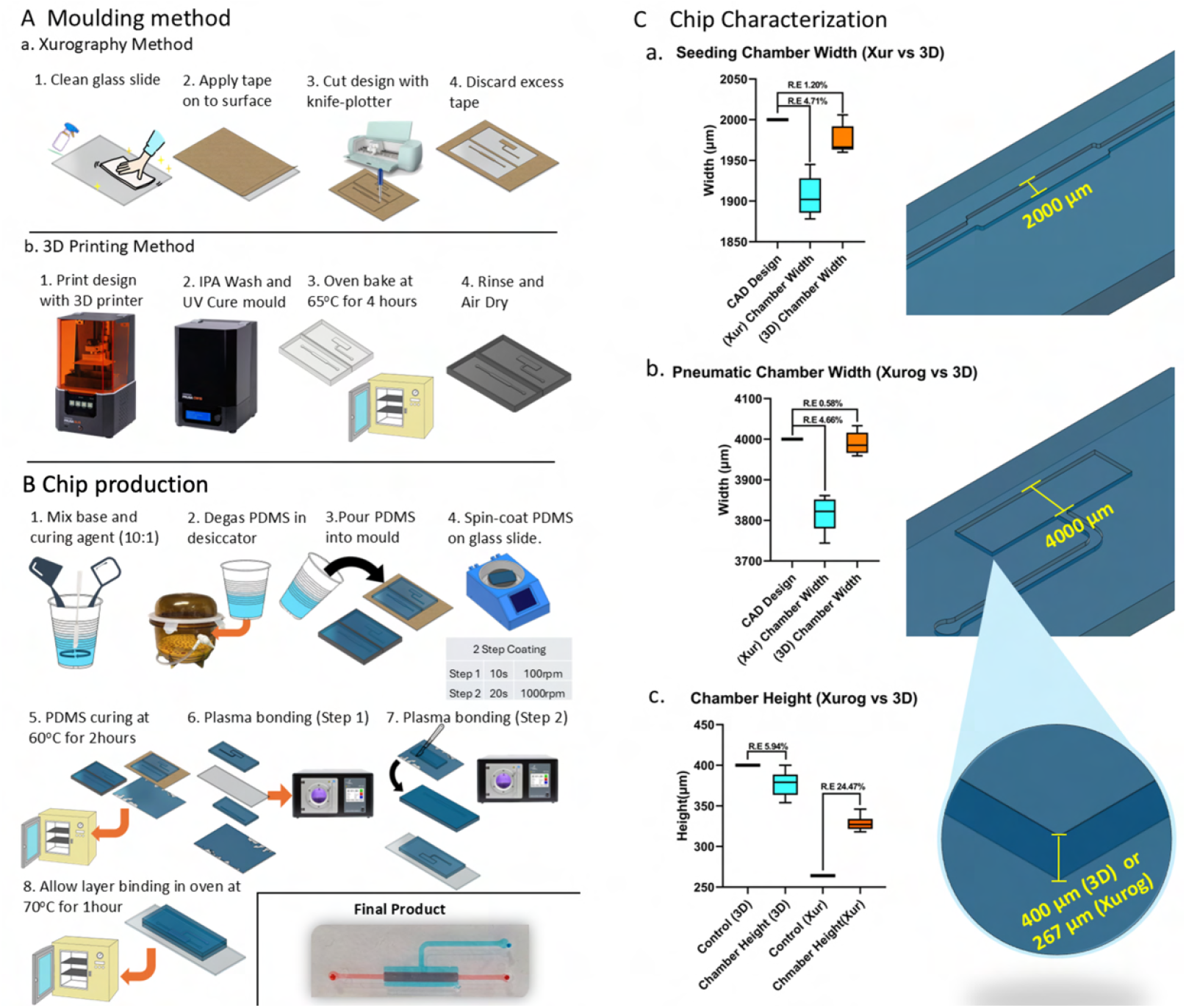
Mould Fabrication Methods and PDMS Chip Production Workflow. The figure illustrates three sections: the first section depicts the xurography method for mould fabrication as Aa, the second section shows the 3D Printing method for mould fabrication A.b, and the final section outlines a common PDMS chip production workflow leading to the final microfluidic chip as B. C: Box plots a, b and c show the difference between the Computational aid design (CAD) design and microscopy measurements, with relative error indicated as “R.E”. Data are presented as mean ± SD, with n=3.

#### 2.1.1 Xurography

The channel designs were plotted in AutoCAD(Autodesk) as a 2D file and exported to Cricut Design Space (Dimensions were shown as FigS1.B). The geometry was selected and sent to Cricut Explore Air 2 (Cricut, UK), a device that employs a knife plotter to cut the design. A glass slide (76×50 mm, Corning) was sequentially cleaned with acetone and isopropanol and then dried on a hot plate at 80°C. The cleaned glass slides were covered with heat-resistant adhesive film (Blackhole lab, France), positioned on the Cricut mat, and inserted into the Cricut machine. The designs were cut, and unwanted regions were excised using forceps, followed by a cleaning process with isopropanol.

#### 2.1.2 3D printing

The mould design was fabricated using the Prusa SL1S printer (Prusa Research a.s., Czech Republic) with Phrozen Aqua-Grey 4k resin (Phrozen Tech Co. Ltd). This printer employed a layer-by-layer UV curing technique to create precise 3D structures. The process began by creating the mould design in AutoCAD (Autodesk), which extrudes the geometry into a 3D shape. The design was printed using the integrated “Ultra Detail” setting with a 25 µm layer height, and the exposure time was set to 1.1 seconds per layer with a 20-second exposure for base layers. The resin was poured into the printer vat after being thoroughly shaken. Printing typically takes less than 30 minutes depending on the design height. Once printing was completed, post-processing was carried out to prepare the mould for PDMS application. The mould was carefully removed from the build plate and washed in isopropanol for 10 minutes to remove uncured resin. After washing, the mould was dried and cured under UV light for 15 minutes using the Prusa CW1S wash and cure station to ensure complete solidification. Following UV curing, the mould was baked in an oven at 60°C for 6 hours. The mould was now ready for PDMS applications. An additional coating process is needed for resins such as the Prusament Tough Black to prevent PDMS from adhering to the mould. A hydrophobic surface treatment was applied by coating the mould with a 1% (v/v) solution of trichloro-perfluorooctyl silane (Sigma-Aldrich, USA) in isopropanol (Fisher Scientific, USA). The coating was allowed to evaporate at 60°C, forming a hydrophobic layer on the mould surface, and ready for PDMS application.

### 2.2 PDMS device fabrication by replica moulding

The PDMS oligomer and its curing agents (SYLGARD 184, Dow Chemicals, USA) were accurately weighed in a 10:1 weight ratio and thoroughly mixed. The solution was then degassed for 30 minutes using a vacuum desiccator to completely remove any trapped air bubbles. The degassed PDMS mixture was carefully poured onto the mould and cured at 70°C for two hours. Post-curing, the PDMS chips were peeled off, inlet and outlet holes were punched, and the chip cleaned by ultrasonication for 2 minutes, and dried using an air gun. A two-step spin coating procedure was employed to prepare the thin PDMS membrane (Step 1: 100 revolutions per minute for 20 seconds, Step 2: 1000 revolutions per minute for 20 seconds). A cleaned glass slide (75×50mm, Corning USA) was positioned on the chuck of the spin coater (Ossila, USA), and 1 ml of the degassed PDMS mixture was carefully pipetted onto the centre of the glass slide, followed by the spin coating process. After spin coating, the glass slides were released and kept at 70°C for 3 hours to complete the curing. The cleaned PDMS chips and membrane were treated with air plasma for 1 minute at 0.48mTorr (Henniker Scientific, Runcorn, UK), bonded by gently pressing each layer together, and kept at 70°C to complete the bonding.

### 2.3 Microfluidic device dimension characterization

Measuring channel dimensions in a PDMS chip involved the use of a Nikon Eclipse NI Upright Fluorescent microscope with a 10x magnification objective lens. To measure the channel length and width, the PDMS chip was placed on the microscope stage, and clear images of the pneumatic and seeding channels were captured. For channel height measurement, the chip was carefully sliced into 2-3 mm thick sections using a scalpel or razor blade. These slices were laid flat on a surface, and the microscope was focused on the cross-section to obtain a clear view of the channel height. Images were taken, and imported into Fiji (ImageJ, NIH, USA) for width, length and height measurements in microns.

### 2.4 Computational simulation

The Solid Mechanics module in COMSOL Multiphysics™ software was used to model the displacement of a PDMS thin membrane between the cellular chamber and the pneumatic chamber under different applied pressures using Finite Element Modeling (FEM). The laminar Flow module was utilized to evaluate the fluid dynamics in the cellular chamber. Briefly, for pneumatic chamber simulation, a 2D design file was employed to reduce the computational load, and the central vertical cross-section of the device was selected for modelling. PDMS was modelled as a liner elastic material (10:1 polymer base/curing agent ratio), which material parameter used in this study showed as follows: ^28^ PDMS density was set to 965 *kg/m*^3^, and the Poisson’s ratio was set to 0.49. The Neo-Hookean model was applied in this study with the Lamé parameter set to 6.6710^5^ *N/m*^2^ and bulk modulus,, of 3.3310^7^*Pa*. Laminar flow physics (SPF) was chosen to simulate fluidic flow through the microfluidic channel, and a stationary study was performed. A ‘no slip’ boundary condition was applied, with the outlet boundary condition defined as outlet pressure (P) = 0. The shear stress was determined by multiplying the shear rate (spf.sr) and the solution viscosity (spf.mu).

### 2.5 Experimental PDMS membrane deflection and linear strain determination

The linear strain exerted due to the pneumatic actuation of the PDMS membrane was calculated by measuring the membrane’s original length and the membrane’s length after stretching along the central axis. To functionalise the membrane, the EOC fluidic channel was briefly treated with 4% (3-Aminopropyl)triethoxysilane (APTES, Sigma-Aldrich) solution in absolute acetone for 20 min, enhancing the binding of fluorescent beads and minimizing non-specific interaction. After incubation, the channels were washed with absolute ethanol and dried at room temperature. Fluorescence beads were (0.5mg/mL,Alexa 647-PS COOH, 0.5µm, Bangs Laboratory) ultrasonicated to get a uniformly dispersed solution. The beads were subsequently introduced to the fluidic channel to visualise the displacement patterns as in ^29^, and the unbounded beads were removed by washing them with absolute ethanol. Later, the chips were dried at room temperature. The pressure source (Elveflow) was connected to the pneumatic actuation channel, applying pressures ranging from −100 to +100 mbar, in 10 mbar increments. The 3D stack images were captured (Nikon Eclipse Ni, Japan), and the membrane length after stretching was calculated from the reconstructed image (Nikon Elements, Nikon, Japan). Three ROIs were selected and measured for each chip, and experiments were conducted separately on three different chips. The linear strain was calculated based on the following formula: ^30^

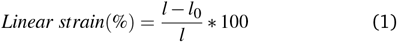

where, l = length of the membrane after expansion and *l*_0_ = Original length.

Using ImageJ, the 3D Object Counter plugin identified bead positions within a defined Region of Interest (ROI), extracting the relevant YYY- and ZZZ-coordinates across the deflecting membrane. The resulting dataset was then imported into OriginPro, where a second-order polynomial curve was fitted to describe bead displacement (vertical axis) in relation to the ROI short-axis position (horizontal axis). It automatically generated both the 95% confidence interval and the 95% prediction band around the fitted curve, providing statistical bounds and a predictive measure of membrane deformation.

### 2.6 Particle Image Velocimetry (PIV) Analysis of PDMS Membrane Deflection

Each image taken from the beads tracking experiment was cropped or selected to define a 512×512 pixel region of interest (ROI) centered on the membrane, thus ensuring consistent coverage across all recorded pressures. In MATLAB’s PIVLab environment, each pair of sequential images was loaded, and the pre-defined 512×512-pixel ROI was selected for uniform comparison. The particle image velocimetry calculations were performed at “Extreme” robustness with 3 pass interrogation area of 128, 64 and 32 pixels. The resulting velocity vectors were scaled to visualize displacement across the membrane surface, and their directions and magnitudes were inspected to assess both localized and global membrane movement. Built-in magnitude analysis tools in PIVLab enabled extraction of displacement values specifically at the center and edges of the membrane, yielding a spatial map of deflectionunder varying pressures. The exported displacement data were subsequently processed to determine average displacements at different pressure increments, generate deflection trends, and evaluate potential spatial heterogeneity in membrane deformation.

### 2.7 Inhaled aerosolised NaCl nanoparticle deposition study

A 1 wt% NaCl in MiliQ water solution was prepared and aerosolised by the Collison 3-jet nebuliser to produce a stable aerosol source. Aerosols were dried in a diffusion dryer (Cambustion) before being size selected and counted by scanning mobility particle sizer (SMPS, TSI SMPS™ 3938), operated with a 10:1 ratio of sheath to inlet flow, as recommended by TSI. A bypass of equal length was used to measure aerosols through the pneumatic chip. The 1*/*4 inch rubber tubing was connected to the 1*/*16 inch chip inlet and outlet through a SwageLok. The chip was pneumatically actuated by TTP Ventus pump (The Lee Company). Each measurement was a single 300-second scan between 12 nm and 300 nm, and each measurement was done in triplicate. A different chip had to be used for each measurement. Penetration was used as the metric for aerosol deposition where:

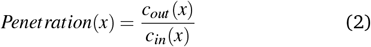

Where *c*_*out*_ (*x*) denotes the outlet number concentration of particle (*x*), and *c*_*in*_(*x*) denotes the inlet number concentration of particle (*x*). Theoretical penetration (*P*) values calculated only for diffusional losses were calculated using equation from Gormley and Kennedy: ^31^

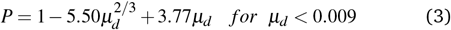

Where:

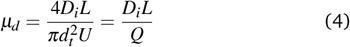

Where *L* is the length of the chip, *Q* is the volumetric flow rate and *D*_*i*_ is the diffusivity, calculated from:

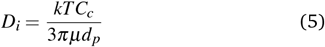

Where *k* is the Boltzmann constant, *µ* is the dynamic viscosity, *d*_*p*_ is particle diameter, and *Cc* is the Cunningham slip correction factor which was calculated using:

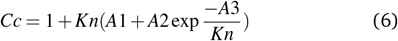

Where *Kn* is the Knudsen number, A1=1.257, A2=0.400, A3=1.100.

A theoretical Stokes number for the system was calculated using the equation:

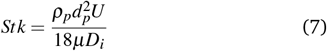

To visualize the spatial deposition within the chip the number of scans was increased to 12. After 12 scans, brightfield imaging of the pneumatic chip was conducted using the 4x Nikon objective lens. An image of the inlet, middle, and outlet of the chip was taken. ImageJ was used to produce a surface plot for the middle image, and histograms of the intensity for each image. The histograms were collated, averaged, and a bar chart of “low intensity” (0-1800) counts was produced where the relative error relates to the spread of deposition within the pneumatic chip.

### 2.8 Cell culture

Human Umbilical Vein Endothelial Cells (HUVEC) were cultured in EBM-2 media (CC-3202, Lonza), prepared as per the manufacturer’s protocol, and maintained at 37°C in a 5% *CO*_2_ incubator. Cells from passages 4 to 9 were utilised for the experiments. The human Alveolar Epithelial Lentivirus-immortalised cells (hAELVi cells) as an alveolar epithelial type I cell line was obtained from InSCREENeX GmbH (Braunschweig, Germany) and cultured in Advanced-DMEM (Thermal Fisher Scientific) supplemented with 1% v/v Newborn Calf Serum (NCS, Sigma-Aldrich) and 1% v/v 1% penicillin/streptomycin(Gibco™). Cells were maintained at 37°C in a 5% *CO*_2_ incubator. Cells from passages 20 to 60 were utilised for the experiments.

### 2.9 PDMS membrane surface coating and optimization of microfluidic cell culture

Before cell seeding on the EOC, channels were sterilised with 70% ethanol (Fisher scientific) rinsed with sterile phosphate-buffered saline (PBS, Sigma-Aldrich), and autoclaved. The sterilised chips designated for hAELVi cells were then coated with a 2 mg/mL 3-hydroxylamine hydrochloride (Dopamine-HCl, Thermo scientific) solution prepared in 10 mM Tris-HCl buffer (pH 8.5, Thermo Fisher Scientific) for 3 hours under room temperature. Dopamine in basic solution will undergo oxidative polymerization and produce polydopamine (PDA) deposited on the PDMS membrane. Then PDA-coated chips were rinsed three times with sterile PBS (Sigma-Aldrich) to remove unbonded PDA, air-dried in the biosafety hood and then sterilised for 30 min via UV irradiation. PDA acts as a bridge that can react with the amine groups from the ECM protein (e.g. Collagen and fibronectin). ^32^ Subsequently, PDA-coated PDMS was filled with 50 µg/ml collagen (Gibco, USA) and 50 µg/ml fibronectin (Sigma-Aldrich) solution diluted in sterile PBS in the 37^°^*C* incubator for 2 hours followed by a wash with sterile PBS and drying. For the HUVEC seeding, the chip was directly coated with a 50µg/mL fibronectin-only solution for 30 minutes without PDA coating, followed by a wash with sterile PBS and drying. HUVEC or hAELVi cells (150000 cells per chip) were introduced into the fluidic channel and allowed for cell attachment. HUVECs were cultured in the EOC chips fabricated using the xurography technique, while hAELVi were cultured in the chips fabricated using 3D printing technology. After cell attachment, the media was replaced to remove any non-adherent cells. The cell viability of the hAELVi and HUVEC cells within the EOC chip was analysed using fluorescent staining. Calcein-AM/PI (TOCRIS) staining was employed to differentiate live and dead cells inside the EOCs. Briefly, the cell culture media inside the chip was replaced with sterile PBS containing Calcein-AM (3 µM) and PI (2.5 µM)and incubated for 30 minutes in the dark. Post-incubaion, the cell channels were washed three times with PBS and imaged using a fluorescent microscope (Nikon Eclipse Ni, Japan). The images were analysed using ImageJ.

### 2.10 Microfluidic cell culture with applied pressure

The pressure controller was initiated after a 12-hour incubation to ensure cell stabilisation in the EOC. The media flow was facilitated through a syringe pump (KD Scientific, USA). In brief, the PTFE tubes (OD: 1*/*16 inch, ID: 1*/*32 inch) were cleaned by pumping with 70% ethanol and dried under UV light for at least 1 hour. The sterilised tubes were connected to the inlet and outlet of the chips and the opening of the pneumatic actuation channel. The media perfusion tubes were connected to the syringe pump via a 21-gauge needle, and the pneumatic channel tube was connected to the pressure controller (OB1 MK4, Elveflow, France).

### 2.11 Validation of barrier function under stretching by immunofluorescence

Immunofluorescence staining was performed to examine the integrity of endothelial junctions under media flow and stretching. The HUVEC cells were fixed using 4% paraformaldehyde at room temperature and rinsed thrice with PBS. As previously described in ^33^, cells were then permeabilised with 0.3% Triton X-100 and blocked using 3% BSA. VE-Cadherin rabbit mAb (Dilution 1:400, Cell Signaling Technology, USA) was added to the chip and incubated overnight at 4°C. The chips were washed with PBS, and Alexa Fluor 488 anti-rabbit IgG (Thermo Fisher Scientific, 10µg/mL) was introduced. Alexa Fluor 647 Phalloidin (0.165µM, Life Technologies) was added to the chip and incubated for 1.5 hours at room temperature, followed by DAPI (1:1000) for 30 minutes. After a final rinse with PBS, images were captured using an upright fluorescent microscope and analysed using ImageJ.

Three days after hAELVi cell seeding on the EOC, cyclic pressure (0 mbar, −30 to +30 mbar, −50 to +50 mbar, −70 to +70 mbar) was applied for 24 hours. The sinusoidal waveform had a frequency of 0.2 Hz and started at its maximum value at the beginning of each cycle. Before immunofluorescence imaging, chips were washed with PBS and then fixed with 4% paraformaldehyde solution (PFA) (Sigma-Aldrich, USA) for 15 min at room temperature. After fixation, cells were washed with PBS for three times and permeabilised with 0.1% Triton X-100 (Sigma-Aldrich, USA) solution under RT for 20min followed by PBS washing. Subsequently, cells were blocked with 1% bovine serum albumin (Sigma-Aldrich, USA) solution for 1 hour at RT. Cells were incubated with Anti-ZO-1 antibody (5µg/ml, Invitrogen, USA) diluted in 0.1% BSA solution for 3 hours at RT in the dark. After washing with PBS, cells were incubated with Phalloidin-Atto 488 (1: 50, Sigma-Aldrich) and counterstained with Hoechst 33342 (1µg/ml, Thermo Fisher Scientific) for 30 min at RT. After the final washing step, the fluidic and pneumatic parts were disassembled, and the middle thin membranes with cells were cut and transferred to the glass slide with the cell facing up. Then, cells were covered by mounting ProLong™ gold antifade mountant (Fisher scientific) and coverslips were added on top for storage imaging. Imaging was obtained by an upright fluorescent microscope (Nikon Eclipse Ni, Japan). Acquired images were processed or quantified, by Fiji (Image J).

### 2.12 Nanoparticle delivery study

After cell seeding on EOC for 24h, hAELVi medium (InSCREENeX, Germany) with nanoparticles(Alexa-647 fluorescent beads : 198nm, 25µg/ml, Bang Laboratory) was infused to the cellular chamber at a rate of 0.5µl/min for 6h under various cyclic stretching conditions(Applied pressure: 0 mbar, −30 to +30 mbar, −50 to +50 mbar, −70 to +70 mbar, all pressure condition varies sinusoidally at 0.2 HZ). After exposure, the cells were washed with PBS, fixed with 4% PFA for 15 minutes, and permeabilized by 0.1% Triton X-100 for 20 minutes. 1% BSA solution was used as the blocking solution. cells were stained with Phalloidin-Atto 488 (Sigma-Aldrich) and Hoechst 33342 (1µg/ml, Thermo Fisher Scientific). After a final rinse with PBS, images were captured using a confocal laser scanning microscope (Leica Microsystems, Germany). Acquired images were processed and the mean fluorescent intensity of the nanoparticle uptake per cell was analysed using ImageJ.

### 2.13 Statistical analysis and software

The statistical analysis was performed using Prism GraphPad software v9, OriginPro and MATLAB. Pictographic depicted on figures were made using biorender. The Kruskal-Wallis Test with corrected Duun’s test was used to analyze data of cell viability and immunofluorescence, which a p-value lower than 0.05 was considered as statistically significant. The One-way ANOVA test with a post-hoc Tukey test was used to analyze the data of nanoparticle exposure on chip, *p < 0.05 *p < 0.05, **p < 0.01, ***p < 0.001 and ****p < 0.0001. All experiments were repeated at least 3 times or more than 3. Data show mean ± standard deviation (SD).

## 3 Results

### 3.1 Moulding method comparison: Xurography and 3D printing

The fabrication of the EOC model involved two steps: Mould fabrication followed by chip fabrication using PDMS. Two rapid, simple, and cost-effective methods were employed for mould fabrication, xurography and 3D printing with detailed procedures shown in Fig.1A. The accuracy of the resulting chips was characterized by comparing the width and height of the cellular and pneumatic channels against the design specifications to calculate relative errors Fig.1C. For width measurements, moulds fabricated using xurography exhibited a higher relative error (above 4%) compared to those made via 3D printing, which had an error of less than 2%. Height measurements showed a more significant difference: xurography moulds demonstrated a large relative error of 24.47%, while 3D-printed moulds had a much smaller error of 5.94%.

### 3.2 Membrane deflection prediction and experimental measurements

The cyclic positive pressure applied to the membrane induced its inflation, enabling cell stretching and expansion. A two-step spin-coating process was used to fabricate a thin PDMS membrane with an average thickness of 150 µm. However, it is important to note that excessive stretching of the membrane may risk rupture, particularly at higher strain levels. To evaluate membrane performance, COMSOL was used to simulate the deflection of the PDMS membrane under constant pressures ranging from −100 mbar to 100 mbar. The simulation results, shown in FigS2.C, illustrate the displacement of the membrane as the applied pressure varied. Displacement increases with pressure, At −10/10 mbar, the membrane exhibited minor deflection around 0.05/0.06 mm, while at −100/100 mbar, the maximum displacement reached approximately 0.24/0.28 mm. The XY displacement field was extracted from bead-tracking images under constant pressure (±30 mBar (orange), ±50 mBar (blue), and ±70 mBar (green) as shown in FigS2.B.An increase in displacement with pressure was observed in both computational predictions and experimental data. However, an unexpected displacement asymmetry was detected, which might be caused by differences in the volume of the pneumatic and fluid chambers. To visualize membrane deflection, fluorescent plastic beads (0.5 µm) were uniformly coated onto a PDMS membrane. Upon pressure application, the attached beads deformed in tandem with the membrane, maintaining their adherence throughout the process. 3D fluorescence imaging was performed under constant pressure to analyze the spatial distribution and movement of the beads in response to membrane deflection as shown in Fig.2D. By tracking and connecting the positions of individual fluorescent beads, the deflection pattern of the membrane was successfully reconstructed (FigS2.A) Under normal physiological conditions, alveolar cells experience a linear strain ranging from approximately 5% to 12%. ^34^ As shown in Fig.2F, the quantification of the membrane’s deflection revealed that the induced linear strain ranged from around 1% to 8%, effectively replicating the strain experienced by healthy alveoli under physiological conditions.

**Fig. 2.**
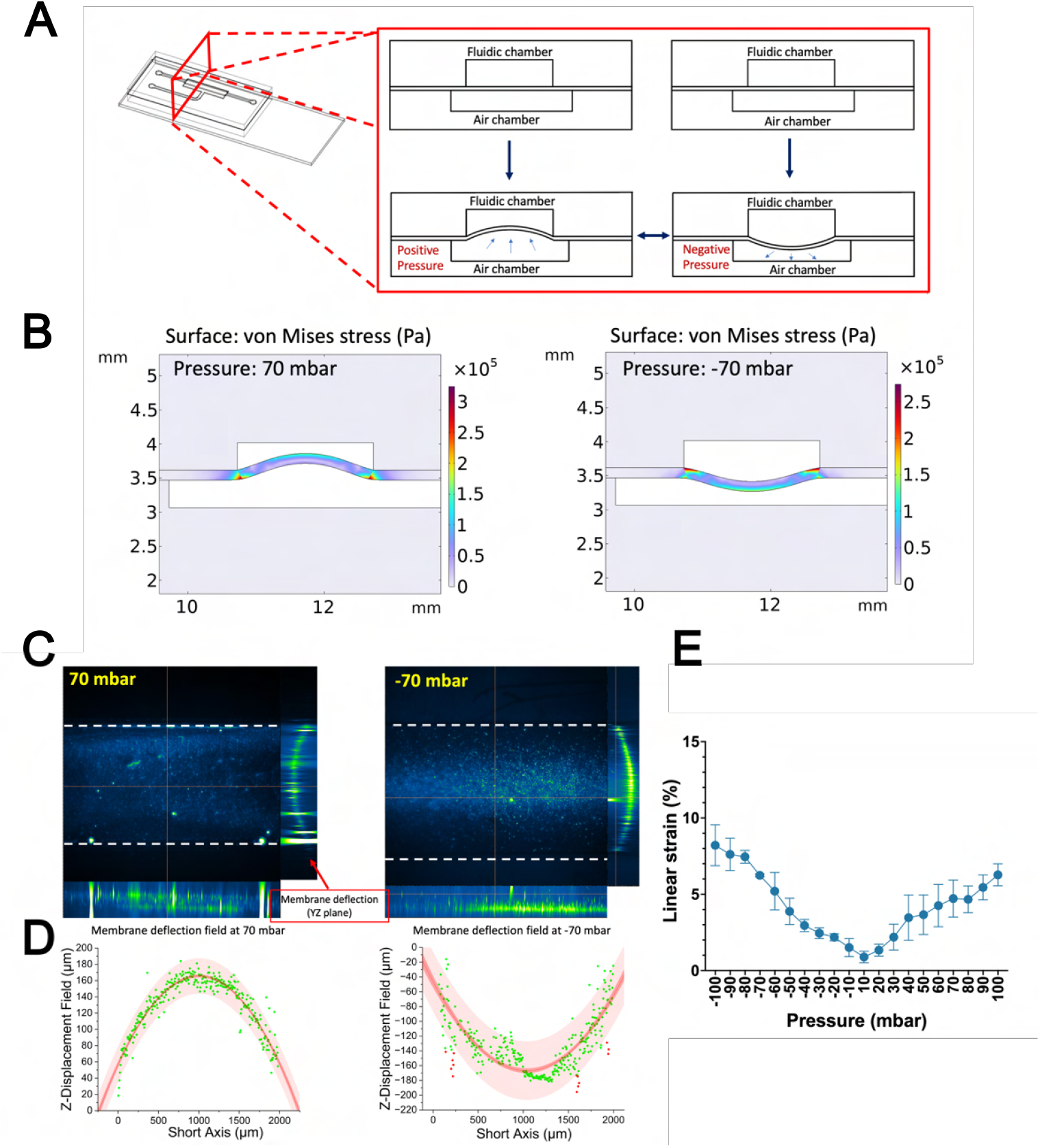
Computational analysis and experimental PDMS membrane deflection applied pressure. (A)Schematic graph of PDMS membrane deflection under positive and negative pressure. (B) COMSOL simulation of the of the von Mises stress (Pa) on the membrane at 70 and −70 mbar, graphs shown as colour heat maps. (C) The cross-section image of the PDMS membrane coated with fluorescent beads at XY-, XZ- and YZ plane was shown. Z-stack images were taken by the fluorescent microscope. (D) A second-order polynomial curve (red line) was fitted to the bead displacement data (green symbols) measured along the short axis of the Region of Interest (ROI). The red-shaded region represents the 95% confidence interval of the fit, while the lighter red band indicates the 95% prediction interval. The vertical axis (in micrometres) corresponds to the measured bead displacement, and the horizontal axis represents the ROI short axis used for the analysis. The fitting is from −70 mbar and 70mbar Bead displacement. (E) The applied pressures and their corresponding linear strains are indicated alongside each image Graph showing the average linear strain under different pressures from −100 to 100 mbar measured from 3 ROI per chip. Data are presented as mean ± SD, with n=3.

### 3.3 Beads displacement field analysis

Fig.3A illustrates displacement vectors for each pressure condition. For positive pressures, vectors point outward, indicating expansion, while for negative pressures, inward-pointing vectors suggest contraction. Displacement increased with pressure magnitude, showing symmetric behaviour across positive and negative pressures, though slight non-linearities appeared at extreme pressures. Particle image velocimetry (PIV) analyses as shown at Fig.3B were conducted to evaluate the displacement of the PDMS membrane under both positive (30 mbar, 50 mbar, 70 mbar) and negative (−30 mbar, −50 mbar, −70 mbar) pressure conditions. Across all pressure magnitudes, a consistent trend was observed: displacement magnitudes increased proportionally with the absolute value of the applied pressure, regardless of its direction. At lower pressures (±30 mbar), displacement maps predominantly exhibited cooler colours (blue), indicating minimal bead movement. As pressure intensified to ±50 mbar and further to ±70 mbar, the colour gradients shifted towards warmer tones (yellow/red), signifying heightened displacement across the membrane surface. The deformation patterns under positive and negative pressures were remarkably similar in both magnitude and directional characteristics. In both scenarios, increasing the absolute pressure resulted in larger displacement magnitudes, and the distinction between central vertical, and peripheral horizontal displacements remained consistent. This bidirectional flexibility underscores the PDMS membrane’s robust mechanical properties, enabling it to respond predictably to pressure variations in either direction. Additionally, localized areas of higher displacement observed as red regions in the colour maps suggest minor inhomogeneities in PDMS thickness or material properties, contributing to spatial non-uniformity in membrane deformation. The displacement analysis revealed distinct spatial variations within the membrane. Under both positive and negative pressures, particles located in the centre of the membrane exhibited greater vertical (out-of-plane) displacement compared to those near the edges, which showed more horizontal (in-plane) movement. These directional trends remained consistent across all pressure levels, suggesting that the central region of the membrane experiences primarily vertical motion, while lateral displacements dominate near the membrane’s periphery. This behaviour likely reflects the mechanical response of the membrane under varying load distributions and boundary conditions.

**Fig. 3.**
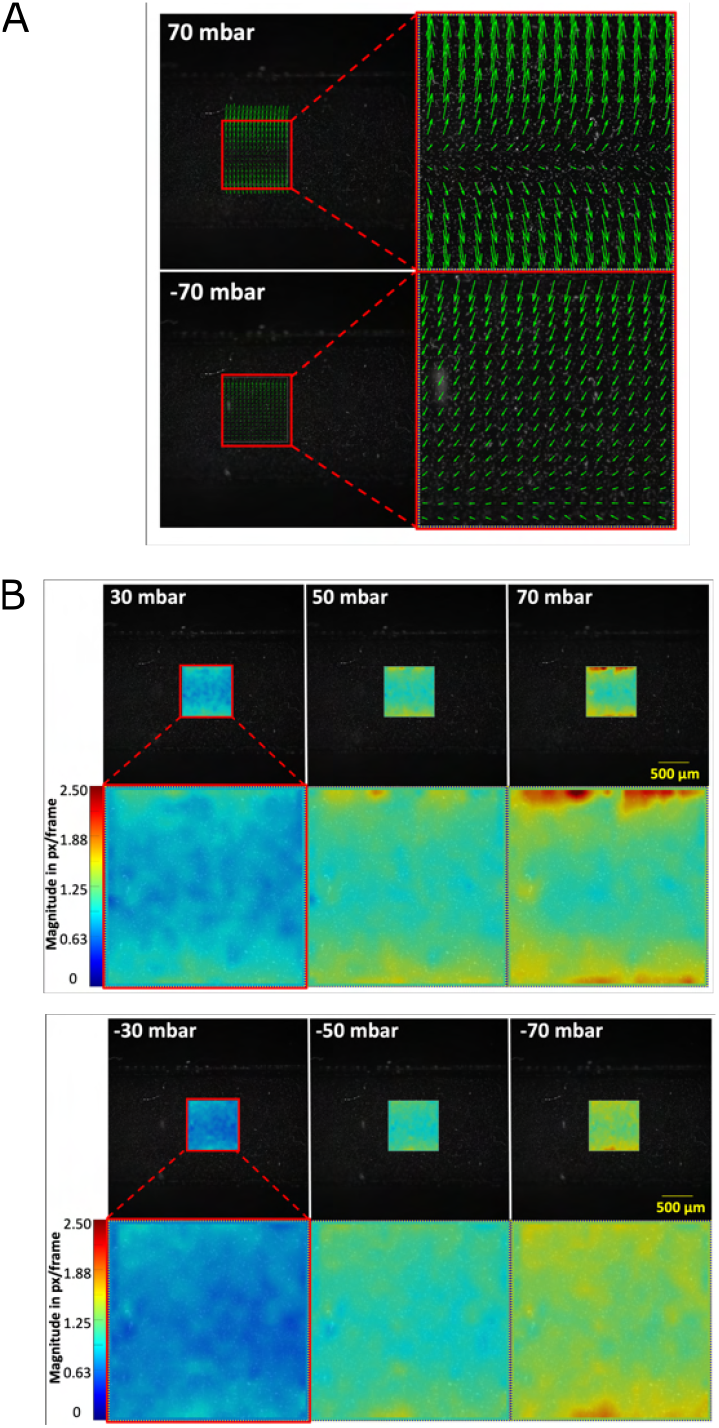
(A)Representative vector field of the PDMS membrane obtained from the PIV analysis of pressures −70 and 70 mBar’s.The arrows indicate local displacement vectors within the 512×512 pixel region of interest, with arrow length scaled to the magnitude of the displacement. (B) Corresponding magnitude map (color-coded) showing the intensity of membrane displacement, with warmer colors representing larger displacements. The color bar provides a quantitative scale for the measured displacement. These visualizations highlight the spatial variation in membrane deflection under applied pressure.

### 3.4 Fluidic Shear stress simulation

Predicting fluid dynamics at the micrometre scale enables simulation of the stresses that cells experience within the EOC model. ^35^ During long-term drug exposure, the cellular chamber is continuously perfused with medium, subjecting cells to sustained fluid shear stress as illustrated in Fig.4A. Therefore, modelling cell drug uptake requires consideration of both shear stress from fluid perfusion and a uniform velocity profile to ensure consistent drug concentration across the cell surface. COMSOL Multiphysics simulations were performed to assess the shear stress and velocity profiles within the fluidic chamber under flow conditions. The simulation results (Fig.4B) demonstrate that fluid velocity is highest in the inlet and outlet chambers, whereas it is lower in the central cellular chamber. This leads to elevated shear stress at the bottom of the inlet and outlet chambers, while the central chamber exhibits lower shear stress. Along the midline of the chamber bottom, the shear stress profile indicates that cells experience maximum shear stress at the centre and minimum near the walls. Moreover, as shown in Fig.4C, shear stress increases with rising fluid velocity. These variations in shear stress may differently affect cell behaviour in various chamber regions. To reduce variability, subsequent studies focused on cells located in the central cellular chamber, where conditions are more uniform., the shear stress experienced by endothelial cells typically ranges from 0.1–5 Pa, depending on the anatomical location, such as veins or large arteries. ^36^ By contrast, the shear stress experienced by alveolar epithelial cells is lower, typically under 1.5 Pa. ^37^ The simulated shear stresses for flow rates ranging from 0.5 to 20 µL/min in this study are all below (Highest shear strain: 0.007Pa) the physiological levels observed *in vivo*. These conditions are predicted to support healthy cell growth in the chip model, even under continuous fluid flow.

**Fig. 4.**
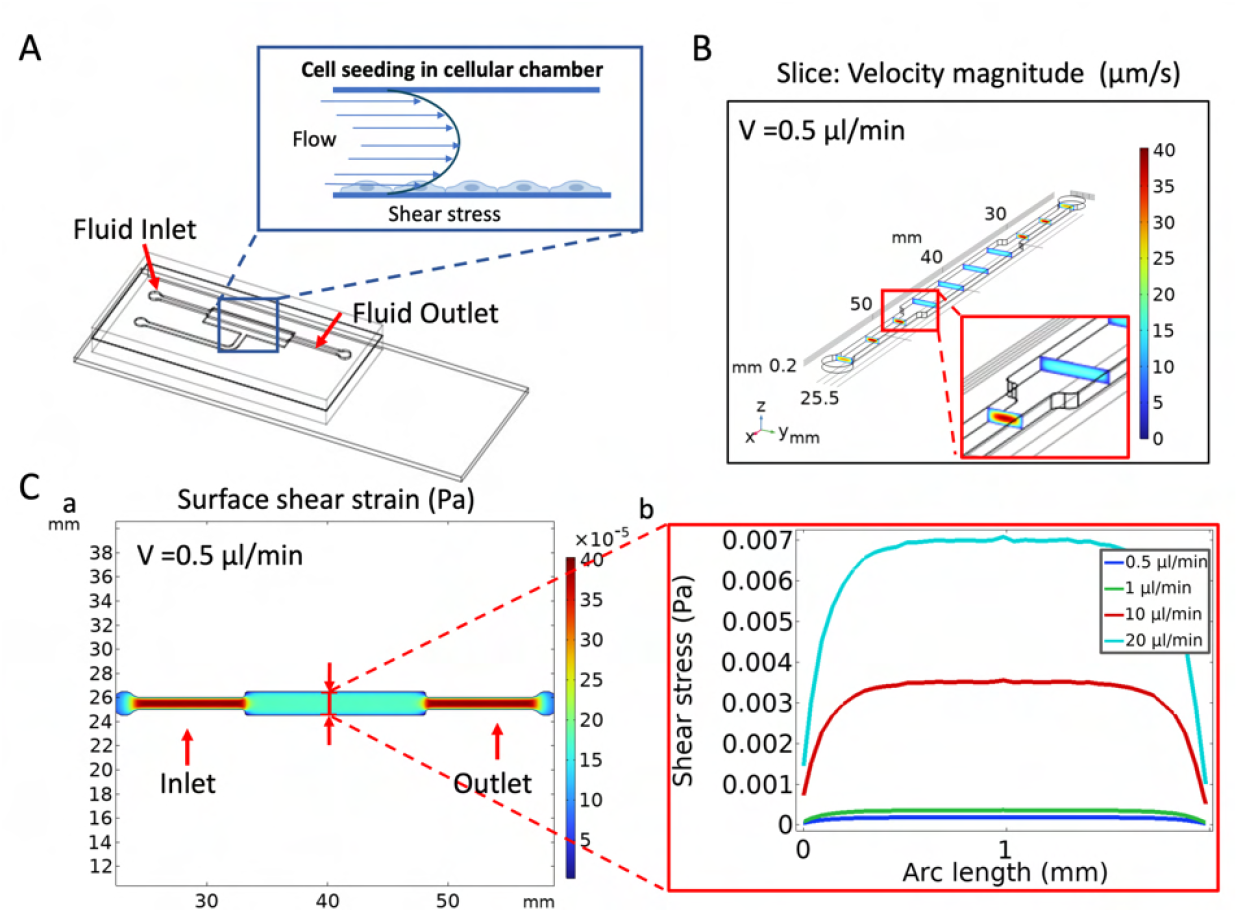
Computational fluid dynamics (CFD) analysis of the shear stress from the fluid to the cells culture on the PDMS membrane integrated to the EOC model. (A) Schematic graph of the EOC model fluidic chamber inlet and outlet. The fluid flow through the chamber will cause shear stress to the cells seeding on the bottom. (B) Predicted velocity magnitude of the media flow in the cellular chamber (Velocity: 0.5µm/min) was performed by COMSOL simulation. Colour map showed higher flow rate at the inlet and outlet channel center, and lower flow rate in the middle chamber center. (C) a) Predicted shear stress surface plot from the media flow (Velocity: 0.5µm/min) to the bottom of the cellular chamber was performed by COMSOL simulation. Colour map showed higher shear pressure at the inlet and outlet channel, and lower uniform shear stress in the chamber for cell seeding. b) The shear stress profile along the chamber bottom midline at different media flow rates (0.5µl/min, 1µl/min, 10µl/min, and 20µl/min).

### 3.5 Inhaled aerosolized NaCl nanoparticle deposition study

The deposition of inhaled nanoparticles in the EOC system under different static deflections was modelled by quantifying the difference in aerosolised NaCl particle number concentration at the inlet and outlet of the EOC system, as illustrated in Fig.5A. The analytical results shown in Fig.5B, show that the deposition is dependent on both the pneumatic pressure applied to the EOC system, as well as the particle size. There is a strong correlation that increasing the membrane deflection increases the deposition within the EOC system. This is corroborated by the imaging results quantified in Fig.5C. The relationship between particle size and deposition is more complex. Nanoparticle deposition within the chip is influenced by impaction, as well as diffusional and electrostatic losses. ^38^ Theoretical calculations of the Stokes number and Penetration (using equations 2-7) for the EOC system are evident against impaction and diffusional losses respectively being the determining factors of deposition in this system. Thus, as also reported by Lidstone-Lane et al, ^39^ it is thought that electrostatic precipitation, caused by the build-up of local electric charge on the non-conductive PDMS walls, is the determining factor in the deposition of the naturally charged NaCl particles. Larger aerosol particles, which have greater surface areas, have a higher probability of carrying multiple elementary charges. For the size range studied here, charge distributions can be described using either the Boltzmann (bidirectional charging), or Fuchs (unidirectional charging) theory. ^40^ Both models predict a significant increase in the occurrence of charge states n = 2,3,4,& 5 for aerosols larger than 50 nm. Higher charge states enhance particle susceptibility to electrostatic deposition due to increased Coulombic interactions. However, larger particles also possess greater inertia, which increases their likelihood of traversing the EOC system. This interplay of forces results in the curve seen in Fig.5B.

**Fig. 5.**
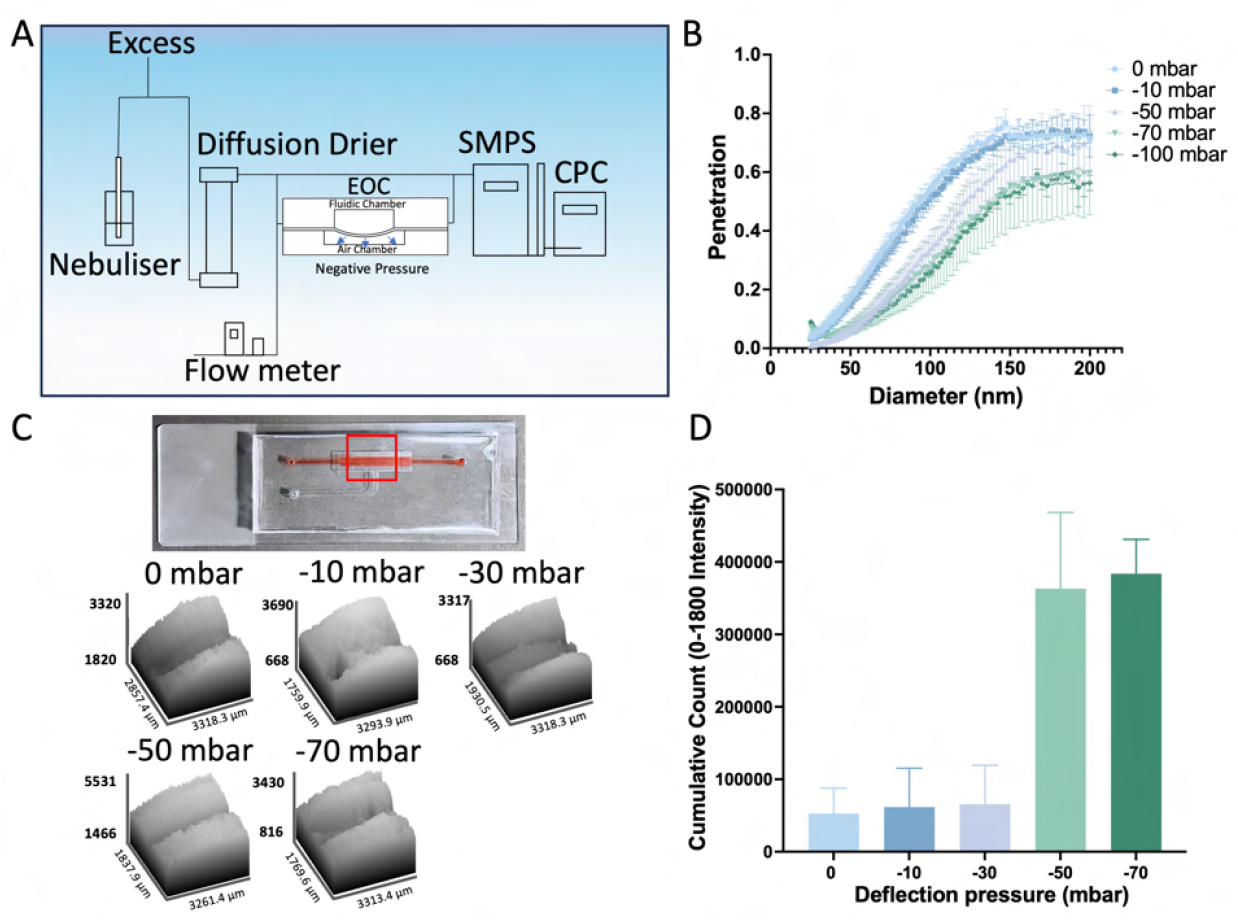
Aerosolised NaCl nanoparticles deposition on chips under constant negative pressure. (A) Experimental set-up for the classification of aerosol deposition through the pneumatic chip. SMPS: scanning mobility particle sizer; CPC: condensation particle counter. (B) Quantification of aerosol deposition at different pneumatic pressures. Penetration was the metric for aerosol deposition, with a penetration value of 1 indicating no deposition. (C) Spatial deposition of aerosols within the chips. (D) Bar chart quantifying the spatial deposition of aerosols within the chip. A 4x objective lens was used to image NaCl deposition at three positions along the membrane to cover its length. A histogram measured the intensity within each image. Through analysis, an intensity threshold of 1800 was identified, indicating the presence of NaCl particles on the membrane. A cumulative count of intensity values ranging from 0 to 1800 was calculated for each image and averaged across the three positions. Data show mean ±SD (n=3).

### 3.6 Effect of applied pressure on endothelial cells growing on EOC

One of the frequent assays employed to evaluate the cell viability in the organ-on-a-chip approach is live/dead assay using calcein-AM/PI dual staining. ^41^ The live/dead assay of static condition (Fig.6A&B) demonstrated that the endothelial cells inside the chips were viable, and they could convert the non-fluorescent calcein-AM dye to fluorescent calcein dye using intracellular esterase enzymes.

**Fig. 6.**
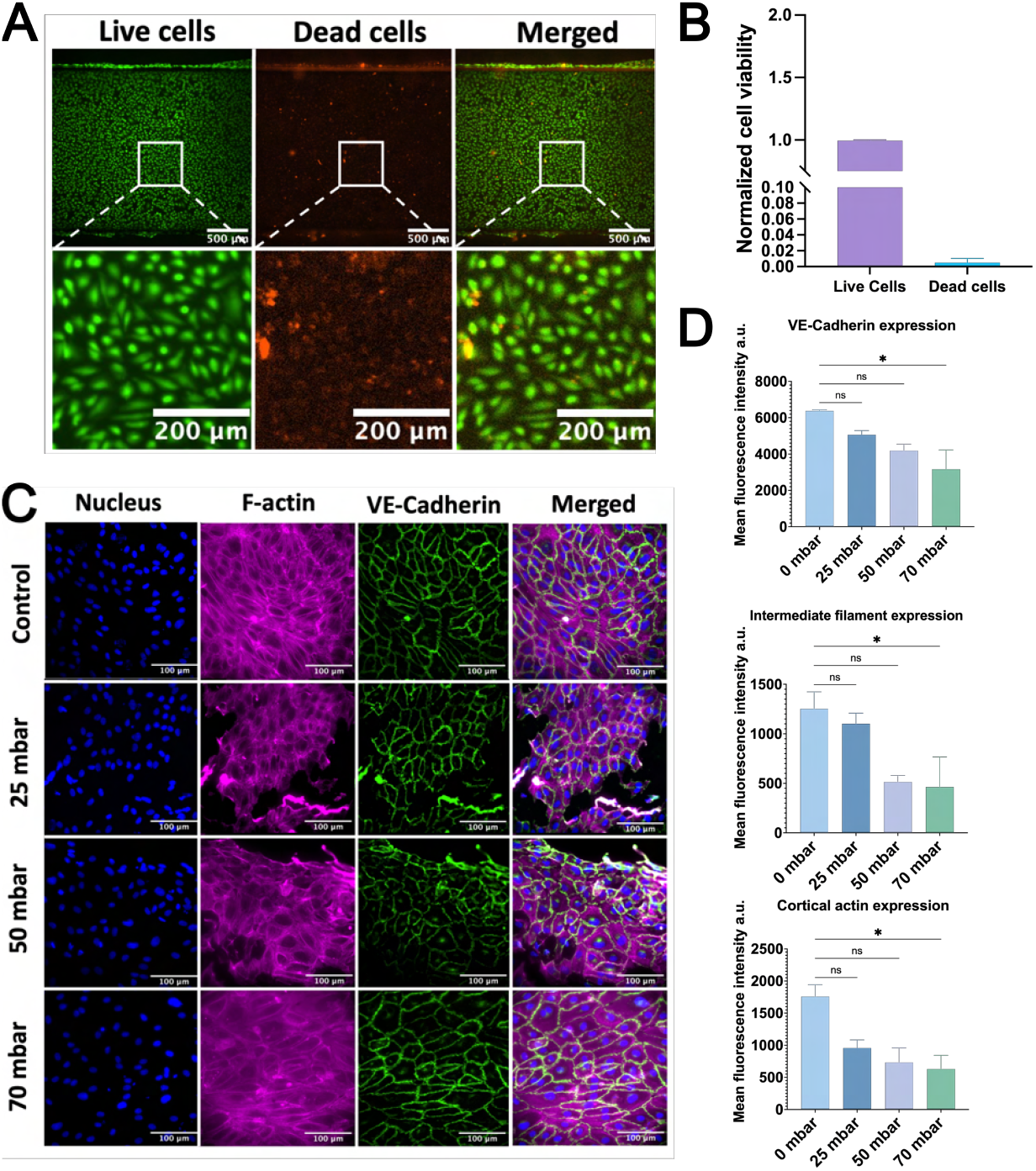
Endothelial cell culture on EOC under constant positive pressure. (A)-(B)Viability of HUVECs on chips. Live/dead fluorescent image of endothelal cells (HUVECs) at static conditions after culturing on fibronectin-coated chips for 24h. Live cells are stained green with calcein-AM, and dead cells are stained red with PI. 99% viability of HUVECs on chips. (C) Protein expression of HUVECs under different conditions, from 0 stretching condition to 70 mbar high stretching pressure. Immunofluorescence image of endothelial cells (HUVECs) at dynamic condition stained for F-Actin Phalloidin (Red), for cell nuclei DAPI (Blue) and for adherens junction VE-cadherin (Green). (D) Mean fluorescent intensity of VE-Cadherins and rhodamine-phalloidin at different stretching conditions. Data analysis is used Kruskal-Wallis Test with corrected Duun’s test *p < 0.05, Data show mean ±SD (n=3).

VE-cadherins are one of the pivotal proteins that control the barrier properties of endothelial cells. ^42^ They take part in cell-cell interaction through adherens junction (AJ). The immunofluorescence assay to visualise VE-cadherins demonstrated intact adherens indicating functionally active AJs. To characterise the impact of stretching on barrier integrity and cytoskeleton integrity of endothelial cells, a range of pressure conditions (0, 25, 50 and 70 mbar) was employed, and the VE-Cadherins and actin filaments were visualized as shown in Fig.6C. The mean fluorescent intensity (MFI) was subsequently calculated (Fig.6D). The mean fluorescent intensity of VE-Cadherins demonstrated a significant stretching pressure-dependent decrease at 70 mbar compare with the control group (0 mbar). This indicates that the integrity of the endothelial cell barrier is compromised as the stretching pressure increases. Similarly, the MFI of both intermediate and cortical filaments also decreased with increasing stretching pressure. Therefore, the stretching pressure impacts both barrier integrity and cytoskeleton integrity. The application of mechanical stretch generally triggers stress fiber formation, which primarily helps in resistance against applied stress mainly in non-muscle cell. ^43^ Further, the thickening of the stress fiber is due to the cell’s adaptation towards the mechanical stress. ^44^

### 3.7 Surface modification of PDMS-based thin membrane with ECM and Polydopamine coating for cell adhesion

Under *in vitro* cell culture conditions, most adherent cell lines require additional support to grow on smooth, hydrophobic surfaces such as glass or PDMS membranes. ^45^ To create a favorable microenvironment for *in vitro* cell growth and proliferation, ECM-mimicking coatings were evaluated. Compared to endothelial cells (HUVEC), hAELVi cells seeding on the EOC experienced more challenges when directly seeding the cells on the hydrophobic PDMS membrane. hAELVi cells were seeded in devices with PDA-coating and without PDA-coating to evaluate the biocompatibility of PDA as an intermediate species to support stronger adherence of collagen and fibronectin coating on smooth PDMS membrane surfaces. hAELVi cell confluency and viability covered on the membrane were quantified as the percentage of total surface area, as shown in Fig.7B. PDA coating on the membrane can significantly improve cell viability on chips from around 25% to 50%, further demonstrating that the PDA-treated method can effectively and biocompatibly serve as a bridge between the ECM and the hydrophobic PDMS membrane without affecting cell growth. In this study, we aim to develop a stretchable thin membrane with high cell adhesion properties. Interestingly, cells growing on chips with a collagen type I coating—one of the most common coatings for adherent cells—did not show high confluency, and we observed the phenomenon of uneven cell distribution (Fig.7A). To test the effect of the ECM on cell growth and proliferation, we seeded cells on chips pre-coated with polydopamine (PDA) and different ECM components: collagen type I, fibronectin, and a mixture of both. Live and dead assay imaging results show that cell viability on the PDMS membrane with the collagen type I-only coating was significantly lower (Around 25%) compared to the mixture (ratio 1:1, Collagen: Fibronectin) coating (Around 50%).

**Fig. 7.**
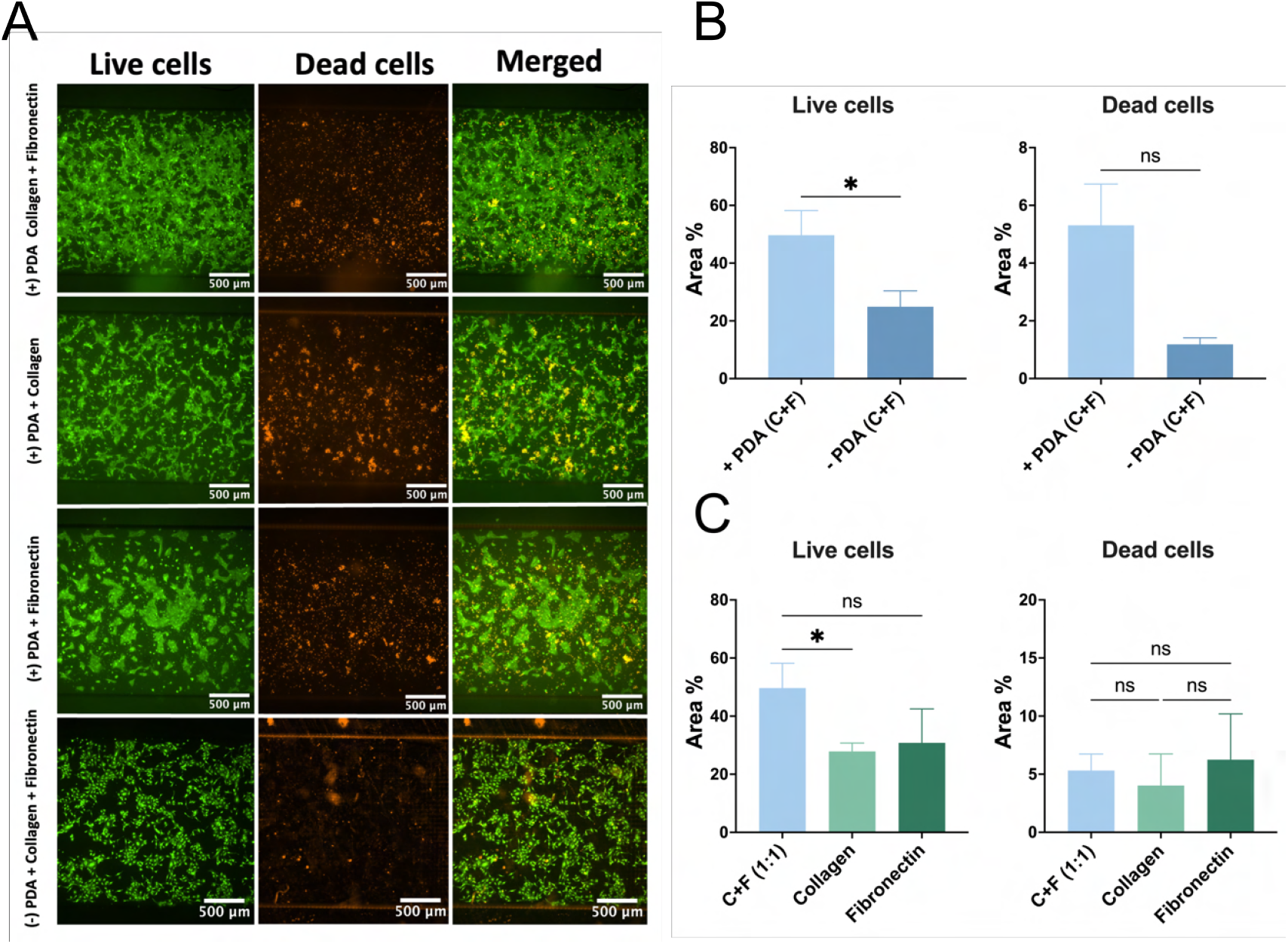
The effect of different coatings of PDMS membrane on the alveolar epithelial cells growing on chips. (A) Live/dead fluorescent image of hAELVi cells growing at PDMS with different coating combination. (B) Viability of hAELVi on in-vitro microfluidic chips analyzed from live/dead fluorescent image of epithelial cells at static conditions after culturing on chips for 48h. Media were changed after 24h cell seeding. Cell viability was compared between PDMS membrane coated with PDA or without PDA, all PDMS membrane were followed with collagen and fibronectin coating (1:1 ratio, 50µg/ml). Live cells are stained green with calcein-AM, and dead cells are stained red with PI. Data analysis was conducted using the Mann-Whitney test, *P < 0.05. Data show mean ±SD (n=4-5). (C) Viability of hAELVi on in-vitro microfluidic chips analyzed from live/dead fluorescent image of epithelial cells at static conditions after culturing on chips for 48h. Media were changed after 24h cell seeding. Cell viability was compared between PDMS membranes were all coated with PDA for 3h. Then, before seeding the cells, the PDMS membrane were coated with collagen(50µg/ml), fibronectin (50µg/ml), Collagen and fibronectin mixtrure (1:1 ratio, 50µg/ml). Live cells are stained green with calcein-AM, and dead cells are stained red with PI. Data analysis is used the Kruskal-Wallis Test with corrected Duun’s test *P < 0.05, Data show mean ±SD (n=3).

### 3.8 Effect of cyclic mechanical strain on alveolar epithelium barrier function

A cyclic stretching of positive and negative pressure on the thin membrane was applied to mimic the human breathing motion *in vivo*. From the live/dead assay result (Fig.8B), there is no significant difference in cell viability between the static control group and the group stretched for 24 hours after 24 hours of cell seeding on chips, but the tendency of decreased cell proliferation can be observed with increased pressures. The barrier function of the alveolar epithelium is regulated by both physiological and pathological signals, with the expression of tight junction proteins serving as a critical determinant. Tight junctions (TJ) are essential for preserving the integrity of the alveolar epithelial barrier by selectively regulating the paracellular transport of molecules and ions between cells and the external environment. ^46^ To further investigate the association between cyclic breathing motion and alveolar epithelium barrier function, we cultured the cells growing on chips for 3 days until TJ protein formed and then applied cyclic pressure for 24 hours. After 4 days of culture under static conditions, the cells were confluent and formed the functional tight junction protein ZO-1 (Fig.8C). The MFI of F-actin expression significantly increased when low pressure (−30 mbar to 30 mbar) was applied, compared to the static condition, as shown in Fig.8D. However, as the pressure increased, F-actin expression did not show any significant differences between the groups. As shown in the Fig.8E, there is no significant difference in the MFI of ZO-1 expression between the static and cyclic stretching groups under different pressures. However, the distribution of MFI of ZO-1 in individual cells shows complexity. The morphology of ZO-1 changed under stretching, as evidenced by the incontinuous ZO-1 barrier. This change in morphology reflects the undulation of the tight junctions.

**Fig. 8.**
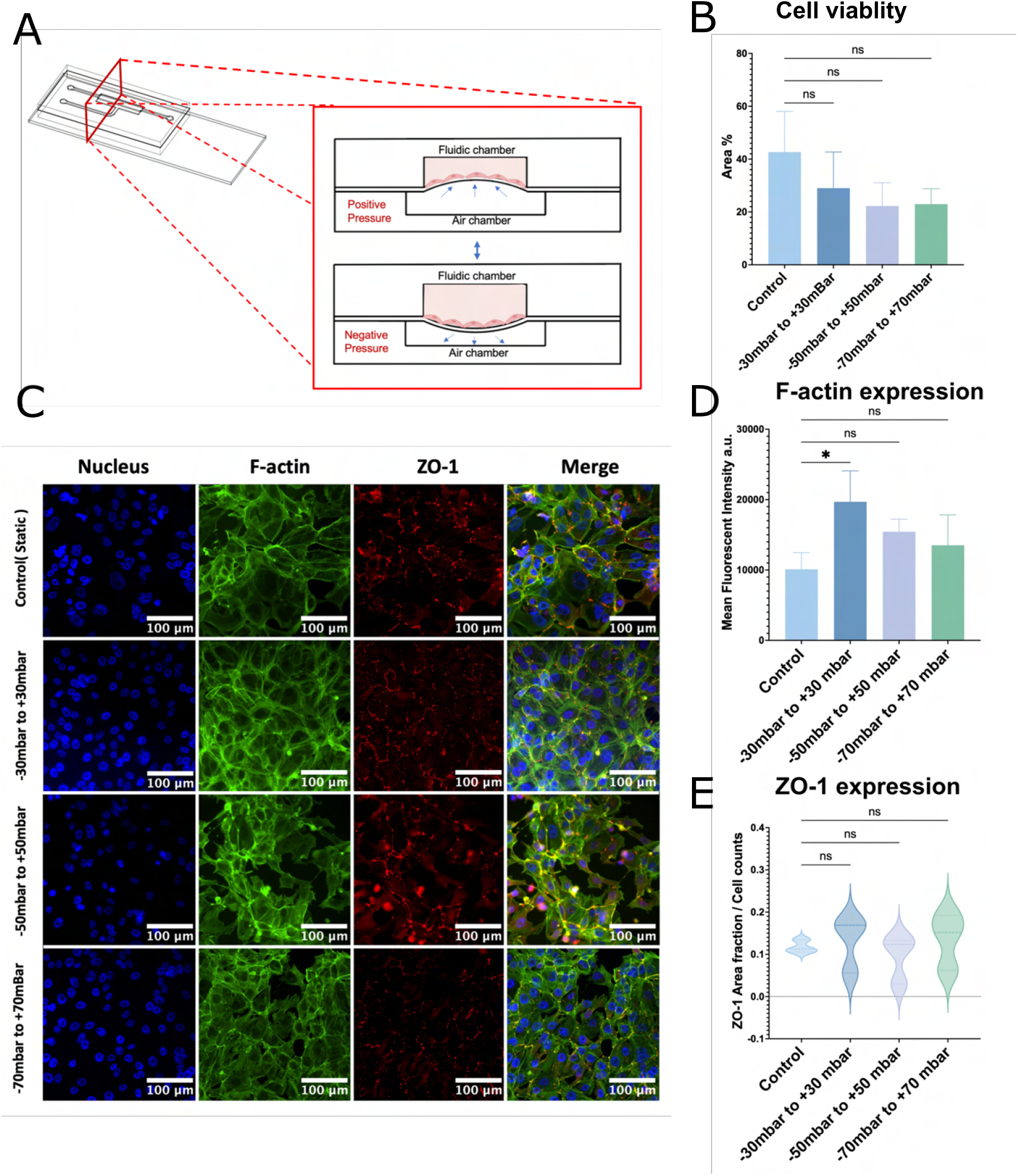
The effect of cyclic stretching on the alveolar epithelial cells growing on chips. (A) Schematic graph of membrane deflection with cells under cyclic stretching. (B) Viability of hAELVi cells on in vitro microfluidic chips analysed from live/dead fluorescent images which were captured after the cells were subjected to stretching for 24 hours. (×20magnification). Media were changed 24 hours post-seeding, and cyclic pressure was applied in the pneumatic chamber to the membrane at −30 to 30 mbar, −50 to 50 mbar, and −70 to 70 mbar. Each pressure cycle lasted 5 seconds with a sinusoidal profile. Data analysis is used Kruskal-Wallis Test with corrected Duun’s test *P < 0.05, Data show mean ±SD (n=3). (C),(D)&(E). Immunofluorescence image of epithelial cells at cyclic stretching condition stained for F-Actin Phalloidin (Green), for cell nuclei Hochest (Blue) and for tight junction ZO-1(Red) (×20magnification). Mean fluorescent intensity of F-actin and mean fluorescent intensity of ZO-1 expression per cell was measured. Data analysis is used the Kruskal-Wallis Test with corrected Duun’s test *p < 0.05, Data show mean ±SD (n=3).

### 3.9 Continuous nanoparticles exposure on chips under cyclic stretching

To better understand the nanoparticle interaction with biological systems, it is necessary to investigate the nanoparticle deposition and intercellular communication particularly in dynamic environments. Cellular uptake of nanoparticles can be influenced by shear stress generated by mechanical forces, such as fluid flow or breathing motions. ^47^ In this study, we examined the cellular uptake of nanoparticles under different pressure conditions over a 6-hour period of continuous nanoparticle exposure. The F-actin and nucleus of the cells were stained to visualise the nanoparticle intracellular uptake as shown in Fig.9A. The results, shown in Fig.9B, reveal that nanoparticle uptake was significantly lower in groups exposed to continuous nanoparticles compared to those with static nanoparticle exposure. Additionally, although no significant differences were observed between the groups, a trend was noted where increased pressure led to a higher accumulation of nanoparticles within the cells. This suggests that while continuous exposure may reduce overall uptake, increased mechanical pressure can facilitate a higher degree of nanoparticle internalization.

**Fig. 9.**
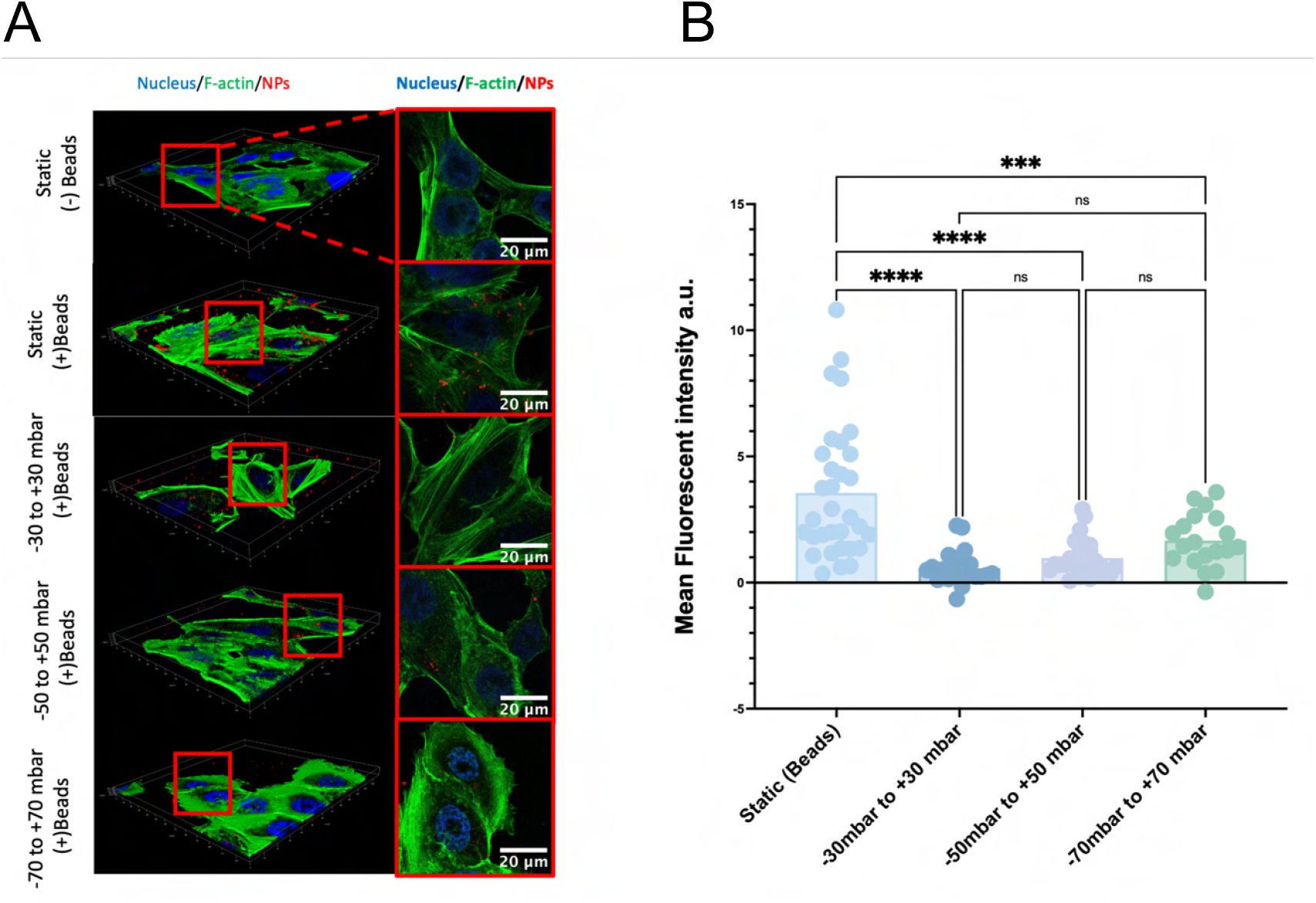
Continuous nanoparticle exposure on alveolar epithelial cells under cyclic stretching for 6 hours. (A) Confocal fluorescent microscopic analysis of cumulative uptake of cells treated with 25µg/ml PS-COOH (Alexa 647, 198nm) diluted in media for 6 h. Images were taken by confocal microscope as Z-stack images (×40 magnification). Cells after treatment were fixed with 4% paraformaldehyde Cell nuclei were stained with Hoechst (blue), F-actin were stained by Phalloidin ATTO 488. (B) Nanoparticle cumulative uptake in individual cells was calculated. Mean fluorescent intensity of the nanoparticle per cell was analysed. Each dot represents one individual cell. Data analysis is used a One-way ANOVA test with a post-hoc Tukey test, *p < 0.05 *p < 0.05, **p < 0.01, ***p < 0.001 and ****p < 0.0001, Data show mean ±SD (n=3).

## 4 Discussion

This study presents an EOC design that successfully replicates the physiological characteristics of alveoli by incorporating breathing-like motions and maintaining continuous, long-term exposure conditions. Using COMSOL simulations, the biomechanical environment within the chip was characterized, effectively mimicking the stretching and fluid dynamics that occur *in vivo* during breathing. This dynamic model not only supports the separate cultures of alveolar epithelial cells (hAELVi) and endothelial cells (HUVECs) on the chip, but also offers valuable insights into the behavior of barrier function under cyclic mechanical stress, which closely reflects the mechanical forces associated with breathing.

A key finding from this study was the critical role of ECM coatings on PDMS membrane in facilitating cell adhesion to the PDMS membrane. While endothelial cells (HUVECs) adhered well to a single fibronectin layer, the alveolar epithelial type I cells (hAELVi cells) required an additional polydopamine (PDA) coating followed with fibronectin and type I collagen mixture coating for stable adhesion. After 48 hours of seeding on the chip, alveolar epithelial cells exhibited higher confluency on a collagen and fibronectin mixture coating compared to single coatings of collagen or fibronectin. A possible reason for this is the deformation of the PDMS membrane, which occurs during subsequent steps such as cell seeding and media changes. Liquid pressure can induce deformation of the restorable PDMS membrane in the microfluidic chips with hollow structures. Even though the PDMS membrane deformation is restorable, the type I collagen fibers coated on the membrane exhibit a non-linear stretching response due to their high stiffness. In contrast, fibronectin fibers are much more compliant and can be stretched to several times their original length. As reported, collagen binding with fibronectin can make fibronectin unfolded and increase flexibility. ^48^ This increased flexibility of fibronectin compared to type I collagen likely contributes to better cell seeding and higher confluency, making the mixture coating more favorable for cell attachment and proliferation. ^49^ Previous works by Michael et.al proved rat alveolar epithelial type II cells growing on fibronectin coating formed a higher-resistance monolayer than collagen or laminin coating in two days, however, ZO-1 expression is lower than the other two coatings. After 5 days cell culturing on fibronectin, the alveolar barrier still maintains subnormal condition compared to cell seeding on collagen or laminin coating. ^50^ This observation prompted us to consider the impact of ECM components on cellular states. While the short-term addition of fibronectin promotes the formation of a monolayer by alveolar epithelial cells, its abnormal accumulation may compromise the integrity of the alveolar epithelial barrier, potentially leading to alveolar fibrosis. Future research should focus on elucidating the effects of ECM components on intracellular signaling pathways to better understand their roles in maintaining cellular homeostasis.

Cyclic stretching was shown to influence epithelial and endothelial barrier properties, underscoring its importance in simulating in vivo conditions. Specifically, stretching reduced the expression of F-actin and adherens junction proteins in HUVECs, indicating weakened cytoskeletal organization and cell-cell adhesion under mechanical stress. This response aligns with previous findings on endothelial sensitivity to mechanical strain, where such stresses affect permeability and barrier function. ^51^ In contrast, hAELVi cells exhibited increased F-actin expression, suggesting enhanced cytoskeletal reorganization and an adaptive response to mechanical forces. Interestingly, the tight junction proteins (ZO-1) in hAELVi cells demonstrated two distinct populations, potentially due to heterogeneity in shear strain distribution across the PDMS membrane. This observation highlights the complex interplay between mechanical strain and epithelial cell behaviour, warranting further investigation into its underlying mechanisms. It has been reported that applied mechanical force can cause ATP depletion, therefore disrupting the ZO-1 structure. ^52^ Under cyclic stretching, the shear strain experienced by cells due to membrane deflection varies depending on the location, with lower strain in the central region and higher strain in the peripheral areas. According to previous research by Chong Shen et al., alveolar epithelial cells cultured on a PDMS membrane in a chip model underwent necrosis when exposed to physiological strain levels (10%), resulting in epithelial damage. ^53^ This suggests that the variation in strain distribution may contribute to localized cell injury, warranting further investigation into its effects on epithelial integrity and inflammatory responses.

The study also evaluated two fabrication methods, xurography and 3D printing, for developing the EOC model. Both methods demonstrated high precision in chip fabrication and supported cellular growth, affirming their suitability for organ-on-chip applications. However, each method has distinct advantages. Xurography, being significantly faster and more flexible, is advantageous for rapid prototyping and iterative design modifications. ^24^ It involves cutting thin films adhered to glass slides, with the channel height determined by the film thickness. However, repeated use of these films can lead to compression over time, resulting in greater variability and significantly larger height errors. In contrast, 3D printing enables the fabrication of intricate three-dimensional microstructures, offering a resolution of 50 µm which used in this study. This high resolution is ideal for complex designs but imposes minor errors in width and height dimensions due to the instrument’s resolution limitations. Both methods proved faster than traditional fabrication techniques, such as photolithography, which require time-intensive steps like photomask preparation and repeated baking processes. ^54^ Therefore, the choice of fabrication method should be guided by specific project requirements, with xurography favouring speed and adaptability and 3D printing suited for high-resolution, complex geometries. The aerosol deposition experiments revealed that for particles exceeding 150 nm in size, deposition efficiency was no longer size-dependent but instead governed by applied pressure. Higher pressures led to greater membrane deflection, enhancing nanoparticle deposition. This pressure-dependent deposition mirrors physiological lung conditions, where increased mechanical forces during breathing facilitate nanoparticle delivery to alveolar surfaces. The same trend was observed in cellular uptake experiments, where higher cyclic pressures resulted in greater nanoparticle internalization by epithelial cells. These findings underscore the critical role of mechanical stretching in influencing particle-cell interactions, with significant implications for inhaled nanomedicine development.

## Conclusions

In this study, we developed a multifunctional Exposure-on-a-Chip (EOC) model designed for dynamic cell culture to mimic the breathing motion of alveoli. The EOC consists of three PDMS-based layers: a fluidic chamber for liquid flow, a pneumatic chamber, and a thin PDMS membrane (150 µm) sandwiched between them. By applying cyclic positive and negative pressure to the pneumatic chamber, the middle membrane undergoes rhythmic upward and downward motion, replicating the expansion and contraction of alveoli during breathing. To fabricate the chip, we utilized soft lithography with both xurography-based moulds and 3D-printed moulds, comparing their efficiency. Both techniques produced chips capable of supporting dynamic cell culture while offering cost-effective and scalable fabrication approaches. Computational simulations using COMSOL Multiphysics were employed to model the fluidic environment and mechanical forces, providing insights into physiological conditions and potential respiratory disease modeling. To enhance cell adhesion and barrier function, the PDMS surface was dual-coated with polydopamine (PDA) and extracellular matrix (ECM)-optimized formulations. Under both static and physiological dynamic conditions, adherent alveolar epithelial cells formed tight monolayers and expressed tight junction proteins, demonstrating the model’s ability to support lung epithelial barrier function. In conclusion, while this EOC model does not incorporate both endothelial and alveolar epithelial cells to fully mimic the alveolar-capillary barrier, it remains a valuable tool for assessing the toxicity and efficacy of inhaled nanomedicines, or the study of inhaled hazardous volatile and airborne compounds, a highly current topic given the societal implications of for instance nanoplastic and vapes ^55,56^. By replicating dynamic breathing motion, it provides a controlled platform to study nanoparticle interactions and pulmonary drug delivery. Investigating the interplay between nanoparticle size, mechanical forces, and cellular uptake under dynamic conditions offers key insights into internalization and retention, contributing to the optimization of inhaled therapies for enhanced efficacy.

## Author contributions

X.L.: visualization, methodology, investigation, formal analysis, data curation, conceptualization, writing – original draft preparation, writing – review & editing. M.E, F.D, : data curation, formal analysis, investigation, validation, visualization, writing – original draft preparation, writing – review & editing. C.Y., T.N.: data curation, formal analysis, investigation, validation, writing – review & editing. C.C., Y.C.: conceptualization, methodology, validation, writing – review & editing. M.S.: methodology, supervision, resources, writing – review & editing. J.X., J.B.d.l.S: conceptualization, supervision, resources, project administration, funding acquisition, writing – review & editing.

## Data availability

All data will be available upon request.

## Acknowledgements

J.B.d.l.S acknowledges support from BBSRC (BB/V019791/1) and the Integrated Biological Imaging Network (IBIN), a Technology Touching Life MRC Network (MR/W024985/1). Also funding from MRC (MR/X013855/1), and the Wellcome Trust (301619/Z/23/Z). M.E and F.D are grateful to the PhD Scholarship program CDT in Aerosol Science supported by the EPSRC. J.X and J.B.d.l.S acknowledge funding from the UK Foreign, Commonwealth and Development Office (FCDO) for the CSC split-site fellowship (INCN 2021-143).

