## Supplementary for "A rapidly prototyped, simple yet versatile dynamic breathing Exposure-on-a-Chip for investigating nanoparticle deposition in the alveoli"

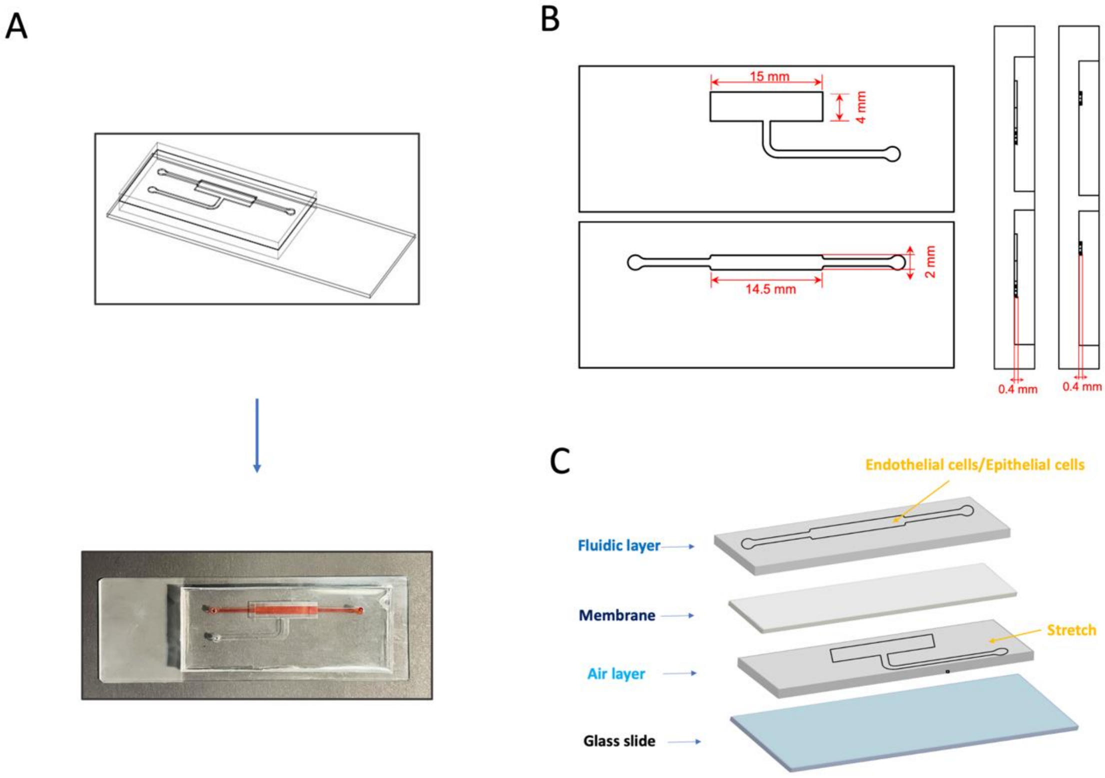


Fig S1: A&B. Schematic design of the Exposure-on-a-Chip device and mould dimensions. C. Graphic illustration of the 3-layered PDMS EOC system. The cellular chamber (top layer), the thin PDMS membrane (middle layer), and the pneumatic chamber (bottom layer).

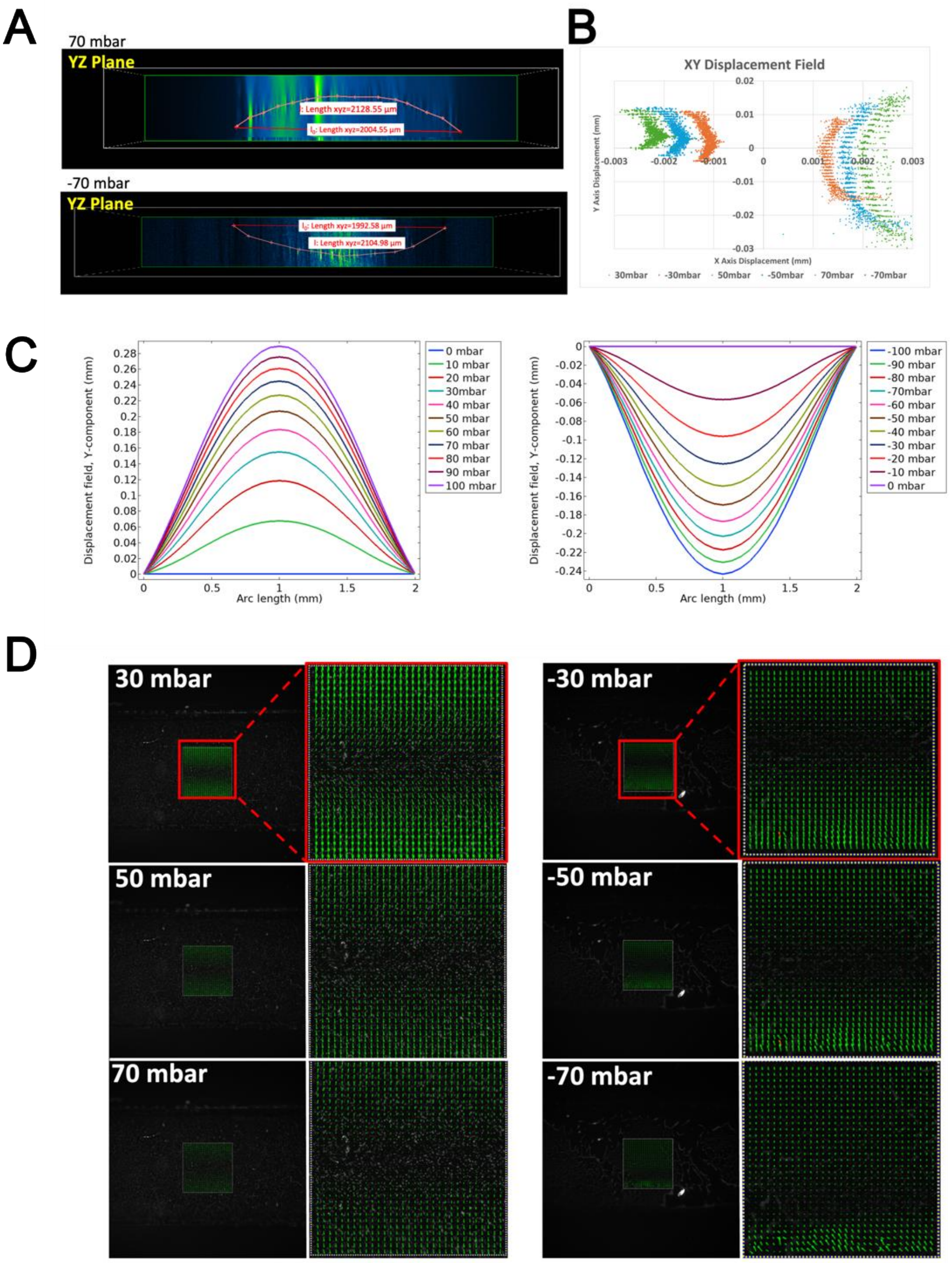


Fig S2: A. Curve of the Beads attached to the membrane are visualized in YZ plane. B. XY displacement field of beads under varying pressure conditions: ±30 mBar (orange), ±50 mBar (blue), and ±70 mBar (green). The plotted points represent bead displacements along the X and Y axes caused by membrane deformation under applied pressures. C. COMSOL simulation of the membrane displacement magnitude at pressure varies from -100 to 100 mbar. D. Representative vector field of the PDMS membrane obtained from the PIV analysis of varying pressures from -70, -50, -30, 30, 50 and 70mBar's. The arrows indicate local displacement vectors within the 512×512 pixel region of interest, with arrow length scaled to the magnitude of the displacement.
